# Multiple beta cell-independent mechanisms drive hypoglycemia in Timothy syndrome

**DOI:** 10.1101/2023.06.16.544987

**Authors:** Maiko Matsui, Lauren E. Lynch, Isabella Distefano, Allison Galante, Nicolas Gómez-Banoy, Hong-Gang Wang, Aravind R. Gade, Daniel S. Sinden, Eric Q. Wei, Adam S. Barnett, Kenneth Johnson, Alfonso Rubio-Navarro, Ang K. Li, Steven O. Marx, Timothy E. McGraw, Paul Thornton, Katherine W. Timothy, James C. Lo, Geoffrey S. Pitt

## Abstract

The canonical G406R gain of function mutation that reduces inactivation and increases Ca^2+^ influx through the *CACNA1C*-encoded Ca_V_1.2 voltage gated Ca^2+^ channel underlies the multisystem disorder Timothy syndrome (TS), characterized by invariant Long QT syndrome and consequent life-threatening arrhythmias. Severe episodic hypoglycemia, which exacerbates arrhythmia risk, is among the myriad non-cardiac TS pathologies that are poorly characterized. While hypoglycemia is thought to result from increased Ca^2+^ influx through Ca_V_1.2 channels in pancreatic beta cells and consequent hyperinsulinism, this mechanism has never been demonstrated due to a lack of informative animal models, thus hampering development of preventive strategies. We generated a Ca_V_1.2 G406R knockin mouse model that recapitulates key TS features including hypoglycemia. Unexpectedly, these mice did not show hyperactive beta cells or hyperinsulinism in the setting of normal intrinsic beta cell function, suggesting dysregulated glucose homeostasis. We discovered multiple alternative contributors to hypoglycemia, including perturbed counterregulatory hormone responses with defects in glucagon secretion and abnormal hypothalamic glucose sensing. Together, these data provide new insights into physiological contributions of the broadly expressed Ca_V_1.2 channel and reveal integrated consequences of the mutant channel that underlie the life-threatening events in TS.

**Brief Summary:** Gain of function mutant Ca_V_1.2 channels drive hypoglycemia through adverse effects on counterregulatory hormones and central nervous system glucose sensing

## Introduction

Almost two decades after the identification of the causal locus and attempts at targeted therapies, the multisystem disorder Timothy syndrome (TS) remains a devastating disease that claims the lives of children. TS results from gain-of-function mutations in *CACNA1C* (1, 2), which encodes the Ca_V_1.2 L- type voltage-gated Ca^2+^ channel. TS mutations. Most common is a glycine to arginine at amino acid 406 (G406R), a homologous residue within either of two alternatively spliced and mutually exclusive exons (exon 8 or exon 8A), that decreases channel inactivation and thereby promotes increased Ca^2+^ influx through the mutant Ca_V_1.2 channel (Ca_V_1.2^TS^) (1, 2). The multiple pathologies associated with TS underline that, compared to other members of the voltage-gated Ca^2+^ channel (VGCC) family, Ca_V_1.2 channels are broadly expressed and contribute to numerous physiological processes (3).

Among the many phenotypes displayed by individuals with TS are life-threatening cardiac arrhythmias exacerbated by sympathetic stimulation and episodic severe hypoglycemia, which also manifests independently (2, 4-7). The pivotal role for Ca_V_1.2 in the cardiac action potential provides a ready rationale for how increased Ca^2+^ influx through the mutant channels prolongs the electrocardiographic QT interval, thus serving as the substate for ventricular arrhythmias (2). Less clear is the etiology for the episodic severe hypoglycemic episodes, present in subjects with either exon 8 or exon 8A G406R mutations (8), thought to trigger as much as 70% of aborted cardiac arrests or sudden deaths in one well characterized TS cohort (9), and suspected of contributing to unsuccessful resuscitations in another (10). This limited understanding presents an obstacle for development of preventive strategies and therapeutic interventions.

The leading hypothesis for TS associated hypoglycemia, although without supporting evidence, is that increased Ca^2+^ influx through the mutant Ca_V_1.2 channels in insulin-secreting pancreatic beta cells leads to inappropriately high insulin secretion (2, 7, 9-12). This hypothesis rests on the role of Ca^2+^ influx through L-type VGCCs to drive the first phase of insulin release from pancreatic beta cells, which express both Ca_V_1.2 and the homologous *CACNA1D*-encoded Ca_V_1.3 L-type Ca^2+^ channels (13). Proposed to support the exaggerated insulin secretion hypothesis is a recent report of a subject with monosymptomatic congenital hyperinsulinism. This subject bears a L566P mutation in *CACNA1C*.

Functional analysis of the mutant channel in a heterologous expression system revealed reduced voltage dependent inactivation analogous to G406R Ca_V_1.2^TS^ channels (12). Yet, since Ca_V_1.2 is the main cardiac Ca^2+^ channel, it is surprising that the subject had neither Long QT syndrome nor cardiac arrhythmias despite an apparent gain-of-function Ca_V_1.2 mutation thought sufficient to drive hyperinsulinism via actions in beta cells. Further, no definitive experimental or clinical data have demonstrated causal hyperinsulinism for any other subject with a Ca_V_1.2 mutation. Thus, whether this mutation is causal for the hyperinsulinism phenotype is not clear. Moreover, the hyperinsulinism hypothesis does not consider the complex regulation of glucose homeostasis that is subject to multiple counterregulatory hormones, secreted by endocrine tissues that also express Ca_V_1.2, nor the growing appreciation for the control of glucose homeostasis by the central nervous system, where Ca_V_1.2 is also broadly expressed.

While investigation of several TS associated phenotypes identified unexpected roles for Ca_V_1.2, mechanistic insight for specific phenotypes, in the context of a broadly expressed mutant has been hampered by the limited availability of animal models. A *Cacna1c^G406R^* knockin mouse model provided insight into autism spectrum disorders associated with TS (14) and separately revealed decreased catecholamine release from isolated adrenal chromaffin cells (15). As catecholamines are a critical regulator in the counterregulatory response to hypoglycemia, those results hint at a potential contributor to hypoglycemia in TS. The *Cacna1c^G406R^* knockin mouse model (“TS2-neo”), however, is a hypomorph, encumbered by the required retention of the neomycin cassette used in gene targeting and a consequent decreased expression of the mutant allele (14, 15). Mouse models that provide opportunities to explore the full scope TS-associated phenotypes, including hypoglycemia, have not yet been reported.

We therefore generated a *Cacna1c^G406R^* knockin mutation in exon 8 by CRISPR/Cas9. This single point mutation avoids the complications of the retained neomycin cassette. The resultant *TS2* mice recapitulate key features of TS, including hypoglycemia, and thus provide a platform to dissect the contributions of various tissues and organs in which the broadly expressed Ca_V_1.2^TS^ channel could drive hypoglycemia. Unexpectedly, we find that hyperinsulinism does not underlie the hypoglycemia in *TS2* mice. Rather, we find that *TS2* mice lack appropriate counterregulatory responses to hypoglycemia.

Further, we identify and confirm expression of Ca_V_1.2 within the arcuate nucleus of the hypothalamus and demonstrate that expression of Ca_V_1.2^TS^ channels in proopiomelanocortin (POMC) expressing neurons is sufficient to promote hyperresponsive glucose sensing. Thus, our data not only provide unexpected insights into how the integrated effects of a widely expressed mutant Ca_V_1.2^TS^ channel lead to TS phenotypes, but more broadly uncover previously unidentified roles for Ca_V_1.2 in physiology.

## Results

The canonical G406 residue mutated in TS resides at the end of the S6 transmembrane segment of domain I in Ca_V_1.2 (**Fig. 1A**). The first two nucleotides of G406 are the last two nucleotides of the mutually exclusive and alternatively spliced exons 8 or 8A, which encode homologous polypeptides with 87% conserved amino acids (**Fig. 1B** - **1C**). Using CRISPR/Cas9, we generated a G406R knockin mutation in exon 8 (*TS2*) (**Fig. 1D**). With exon specific primers, we queried the expression of exon 8 and exon 8a by RT-qPCR in the cerebral cortex, where *Cacna1c* is broadly expressed (16), and observed no exon-specific changes or compensation in expression in heterozygous *TS2* mice compared to *WT* controls, suggesting that the point mutation does not affect expression from the mutant allele (**Fig. 1E**) and thus circumventing the reduced expression of the mutant allele observed in the TS2-neo mice.

**Figure 1:**
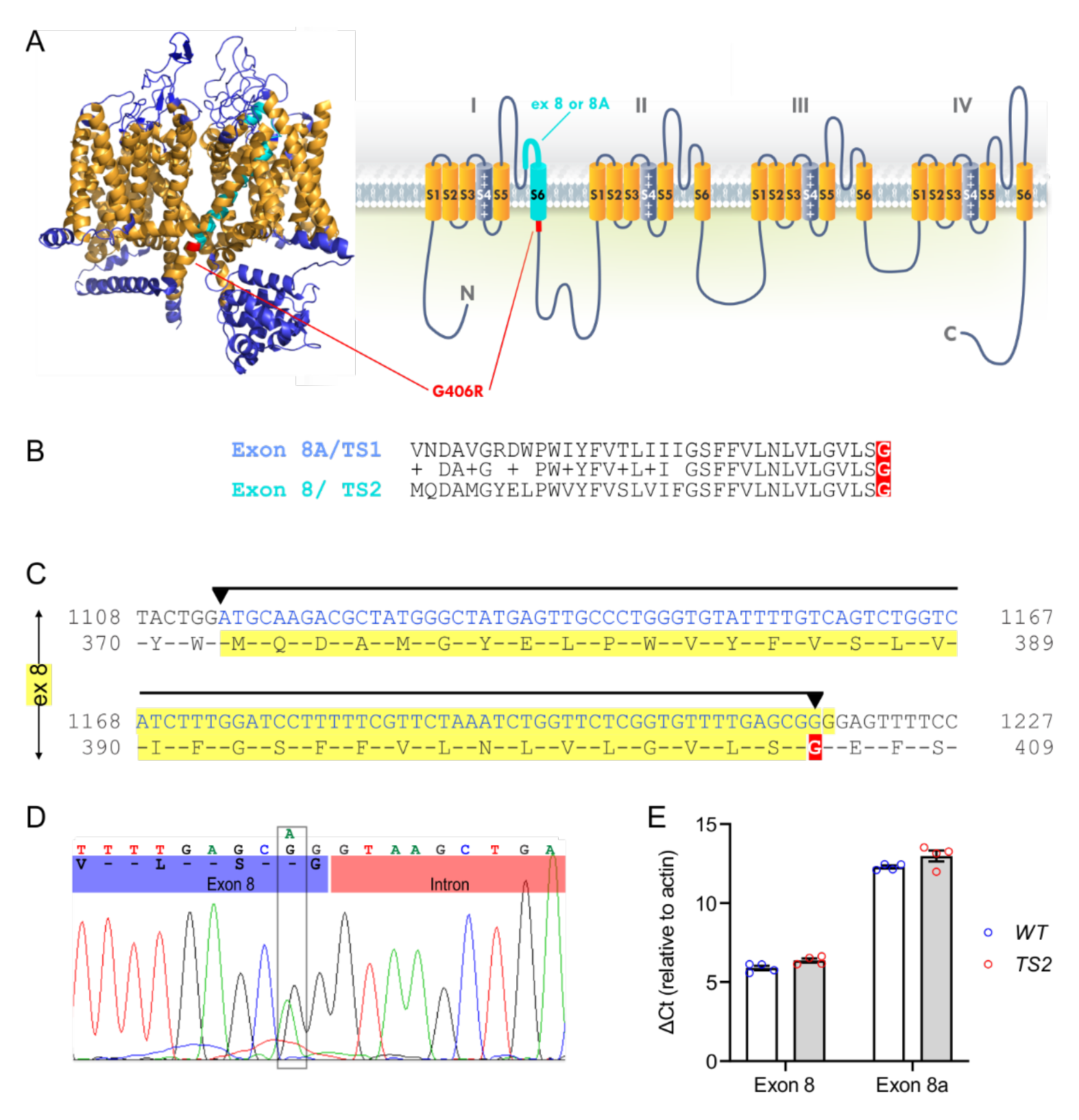
Generation of *TS2* mouse. *A*, Left: structure of the homologous Ca_V_1.1 channel (PDB: 5GJV) viewed from the membrane. The residue homologous to G406 in Ca_V_1.2 is highlighted red. Right: schematic of the pore-forming α_1C_ subunit of Ca_V_1.2. The alternatively spliced exons 8 or exon 8A-encoded peptide that includes the IS6 transmembrane segment is highlighted in light blue and G406 are highlighted in red. *B*, Comparison between the peptides encoded by exon 8 or exon 8a, with G406 highlighted in red. *C*, A portion of the Ca_V_1.2 cDNA sequence containing exon 8 (5’ and 3’ boundaries indicated by arrow heads), with G406 highlighted in red. *D*, Genomic sequencing from a *TS2* mouse showing the single point mutation for the G406R mutation in one of the alleles. *E*, Relative ΔCt values for exon 8 and exon 8a from qPCR using RNA isolated from cerebral cortex of *WT* or *TS2* mice.

Genotyping at weaning showed no homozygous *TS2* mice from more than 20 breeding pairs of *TS2* heterozygous mice suggesting embryonic lethality, consistent with the absence of known homozygous human subjects. Moreover, from 525 pups from *TS2* heterozygous (*Cacna1c^+/G406R^*) × *WT* matings, genotyping at weaning showed reduced numbers of both male and female *TS2* heterozygous mice (99 and 117, respectively) compared to the expected distribution (*χ^2^*=18.4, *p*<0.001). The reason(s) for reduced survival of *TS2* heterozygous mice are not known but may relate to failure to thrive as the *TS2* mice were significantly smaller at weaning (and remained so into adulthood), and the weights of *TS2* male and *TS2* female mice were significantly lower at 10-12 weeks of age (**Fig. S1A-B**). Consistent with reduced weight, in a metabolic cage experiment, *TS2* males ate less that WT males (**Fig. S1C**). Hereon, *TS2* refers to heterozygous *Cacna1c^TS2/WT^* mice.

To confirm the mutant allele was expressed in *TS2* mice, we queried for the Ca_V_1.2^TS^ channel’s signature defective voltage-dependent inactivation (1, 2) in cells known to express Ca_V_1.2. For example, we isolated hippocampal neurons from *WT* or *TS2* postnatal pups (P2) and we analyzed the kinetics of VGCC currents using Ba^2+^ as the charge carrier, thus eliminating confounding effects of Ca^2+^ dependent inactivation inherent to Ca_V_1.2 channels (17). Voltage steps from a holding potential of −60 mV to 0 mV revealed the expected slow voltage-dependent inactivation in *TS2* neurons (**Fig. S1D-E**), significantly slower than the inactivation seen in currents recorded in neurons isolated from *WT* littermates. Since this protocol did not attempt to isolate the L-type Ca^2+^ channel currents from other voltage gated Ca^2+^ channels present in neurons, the Ca_V_1.2^TS^-specific effects on inactivation are diluted, and thus even larger than observed here. We also examined Ca^2+^ channel currents from smooth muscle cells isolated from the colon, as Ca_V_1.2 is the major VGCC in smooth muscle (18). Similar to results in neurons, when using Ba^2+^ as a charge carrier VGCC currents from *TS2* colon smooth muscle cells revealed a marked reduction in inactivation (**Fig. S1F-G**). Thus, VGCC currents in two different tissues demonstrated the significant functional expression of the Ca_V_1.2^TS^ channel. In the adult mouse heart, exon 8-containing *Cacna1c* transcripts dominate, so we expected a prolonged electrocardiographic rate-corrected QT (QTc) interval in the *TS2* mice. Indeed, the baseline QTc interval was prolonged in *TS2* vs. *WT* mice (52.3±1.2 ms vs. 46.3±1.7 ms; *N*=3. *p*=<0.05). Since arrhythmias are exacerbated in TS by sympathetic stimulation (4-6), in a separate experiment we examined the QTc after injection of the non-selective β-adrenergic agonist isoproterenol (1.5 mg/kg) intraperitoneally (i.p.) and observed that at 15 sec after isoproterenol administration the QTc in *TS2* prolonged and remained prolonged 7 min later (**Fig. S1H-I**). Thus, not only are the Ca_V_1.2^TS^ channels expressed in *TS2* mice, but their expression recapitulates the canonical prolonged electrographic QTc interval and promotes arrhythmias after adrenergic stimulation. In line with a recent systematic analysis that noted hypoglycemia in TS subjects with either exon 8 or exon 8A mutations (8), this *TS2* (exon 8) model thus offers an opportunity to investigate the mechanisms underlying the episodic severe hypoglycemic episodes in TS patients.

Several of the reported hypoglycemic episodes in TS subjects occurred after prolonged fasts, so we first measured blood glucose during an extended fast (**Fig. 2A**). For this experiment, we studied mice 8-10 weeks old, as most hypoglycemic episodes in TS patients have occurred in younger individuals (data collected by K.T.). In the fed state (*t*=0), we detected a trend towards reduced blood glucose in *TS2* mice. By 2 h of fasting, blood glucose in the *TS2* mice was lower than in *WT* and remained lower for the entire testing period. These data suggest that the *TS2* mice recapitulate the hypoglycemia phenotype observed in TS. Assess glucose homeostasis, we first tested the response to a 2 g/kg glucose bolus i.p. given after a 16 h fast (glucose tolerance test [GTT]). For this and all subsequent experiments (unless indicated), we used a slightly older cohort (14-20 week old mice), as—consistent with hypoglycemia observed during the prolonged fast (**Fig. 2A**)—the 8-10 week old mice were sensitive not only to the standard prolonged (16 h, **Fig. S2**) fast before a GTT, but the young mice were also unable to survive other standard metabolic challenges, such as an insulin tolerance test (ITT, see e.g., **Fig. 6A**). Even when we halved the standard insulin administration for an ITT, the 8-10 week old TS mice died within 45 minutes of insulin administration. The older cohort, however, survived ITTs with a reduced insulin administration protocol (see Methods), allowing us to query various metabolic parameters. In the 10-14 week old mice, baseline blood glucose before the bolus (*t*=0) was not different between *TS2* and *WT*, but the *TS2* mice showed a reduced glucose excursion compared to the *WT* (**Fig. 2B**).

**Figure 2:**
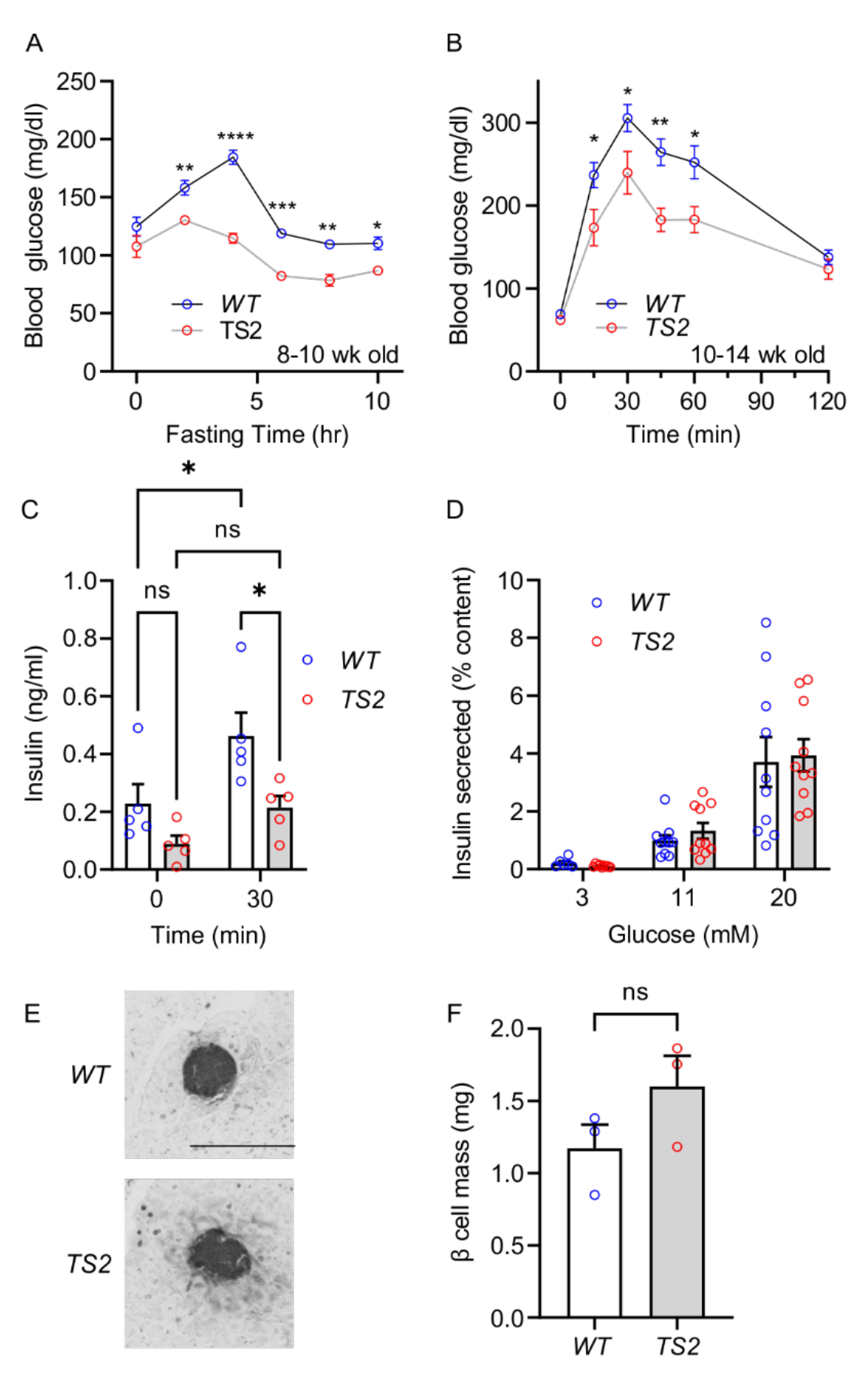
*TS2* mice show hypoglycemia and increased glucose tolerance independent of beta cell insulin secretion. *A*, Blood glucose over a 10 h fast (n=6 each; ANOVA corrected for multiple comparisons; *, *p*<0.05; **, *p*<0.01; and ***, *p*<0.001). *B*, GTT (n=6 each; ANOVA corrected for multiple comparisons; *, *p*<0.05; and **, *p*<0.01). *C*, Basal serum insulin (0 min) and 30 min after i.p. glucose administration (n=5 each; ANOVA corrected for multiple comparisons; *, *p*<0.05). *D*, Glucose stimulated insulin secretion from isolated islets (n=6-10). *E*, Exemplar islet insulin immunohistochemistry. Scale bar, 200 µm. *F*, Pancreatic beta cell mass (n=3 mice/genotype; n=3-4 sections/mouse).

As L-type Ca^2+^ channels are a primary driver of the first phase of insulin release (19, 20), increased insulin secretion driven by Ca_V_1.2^TS^ gain-of-function mutation channels expressed in pancreatic beta cells is the leading hypothesis for episodic severe hypoglycemia in TS patients (2, 7, 9-11, 13). When we measured serum insulin, however, we unexpectedly found that insulin levels were lower in *TS2* compared to *WT* mice in both the fasted state and 30 min after a glucose bolus. Both the *WT* and *TS2* mice showed an increase in serum insulin after glucose administration, but the response was markedly blunted rather than exaggerated in *TS2* mice (**Fig. 2C**). Since the TS2 mice had lower blood glucose levels at 30 min, we therefore assessed intrinsic beta cell function in vitro where we can control glucose levels in glucose-stimulated insulin secretion (GSIS) assays with isolated pancreatic islets. At three different levels of glucose, we observed no differences in the amount of insulin secreted between genotypes (**Fig. 2D**). Further, we calculated beta cell mass and observed no difference between genotypes (**Fig. 2E-F**). The discrepancy between serum insulin (suppressed in *TS2*) and GSIS (similar between genotypes), suggests that in these Ca_V_1.2^TS^ mice insulin secretion from pancreatic beta cells *in vivo* is subject to additional regulatory controls.

In the context of low serum insulin, we hypothesized that the reduced body weight of the *TS2* mice reflected decreased fat storage. Indeed, the major adipose depots (visceral and subcutaneous) were markedly reduced in the *TS2* mice compared to *WT* (**Fig. 3A-B**). Consistent with reduced adipose depots, serum leptin levels were diminished in *TS2* compared to *WT* (**Fig. 3C**). Also consistent with hypoinsulinemia, *TS2* mice showed elevated blood ketones in the fasted state (**Fig. 3D**). Intriguingly, a recent clinical review of TS hypoglycemic cases reported presence of ketones in some patients (12). Together, these data are consistent with hypoinsulinemia, rather than hyperinsulinism in TS.

**Figure 3:**
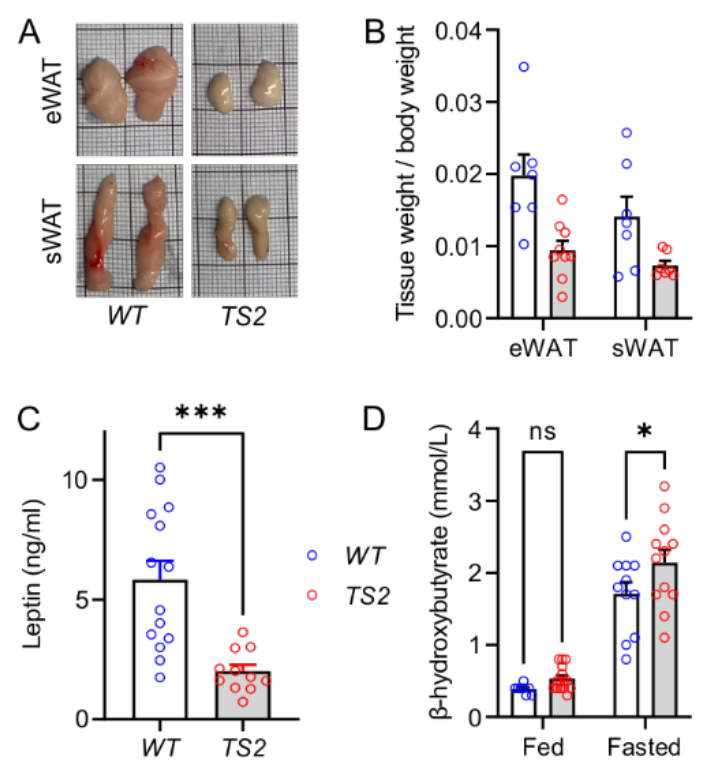
Decreased white adipose depots and increased ketones are consistent with a hypoinsulinemic state in *TS2* mice. *A*, Exemplar epididymal and subcutaneous fat depots (eWAT and sWAT, respectively) isolated from *WT* and *TS2* mice. *B*, Fat depot weight normalized to body weight (n=7-9, ANOVA, corrected for multiple comparisons, *, *p*<0.05; ***, *p*<0.001). *C*, Serum leptin (n=11-14, unpaired *t*-test, ***, *p*<0.001). *D*, Serum β-hydroxybutyrate in fed and fasted states (n=9-15, ANOVA corrected for multiple comparisons, *, *p*<0.05).

Because hypoinsulinemia was unexpected, we extended our investigations to probe the consequences of expressing Ca_V_1.2^TS^ specifically in beta cells. For this, we exploited a mouse model in which a STOP-floxed TS mutant Ca_V_1.2 (Ca_V_1.2^TS^) cDNA has been knocked into the *Rosa26* locus (21) and we activated the transgene specifically in pancreatic beta cells by crossing these mice to a line that expressed a tamoxifen inducible, beta cell-restricted *Cre* recombinase driven by the mouse insulin promoter (22) (**Fig. 4A**). We note that this strategy drives expression of a G406R mutant Ca_V_1.2 cDNA and is thus independent of any differential expression of the mutually exclusive alternatively spliced exons 8 or 8A from genomic DNA that occurs in at least some tissues (23). Compared to non-transgenic controls, in a GTT we observed no differences in basal blood glucose (*t=*0) nor after an i.p. glucose bolus (**Fig. 4B**). We generated an additional mouse line using an alternative, constitutively active *Cre* under control of rat insulin promoter (24), which also failed to elicit a differential response in a GTT (**Fig. S3A**). Although we previously validated that this *Cre*-dependent Ca_V_1.2^TS^ transgene strategy using several different *Cre* lines (25-27), we further confirmed its expression here by employing a cardiomyocyte-specific tamoxifen-inducible *Cre* (*MerCreMer*) (28) to drive expression of Ca_V_1.2^TS^ in cardiomyocytes. In the resulting offspring we observed a profound tamoxifen-dependent increase in the electrocardiographic rate-corrected QT interval (QTc) (**Fig. S4B-C**). (30, 31)Together, these data suggest that expression of a Ca_V_1.2^TS^ in pancreatic beta cells, whether the mutation resides in exon 8 or 8a, does not generate increased insulin secretion. (32)Supporting that hypothesis, we queried *Cacna1c* and *Cacna1d* expression from a single cell RNA-seq data set (32), which showed that *Cacna1c* is the minor L-type Ca_V_1.2 Ca^2+^ channel in beta cells (**Fig. 4C**). Separately, analysis of a human single cell RNA-seq data set (33) also showed that *CACNA1C* is the minor L-type Ca_V_1.2 Ca^2+^ channel not only in beta cells, but within other islet cells. We therefore considered alternative mechanisms for hypoglycemia in the *TS2* mice.

**Figure 4:**
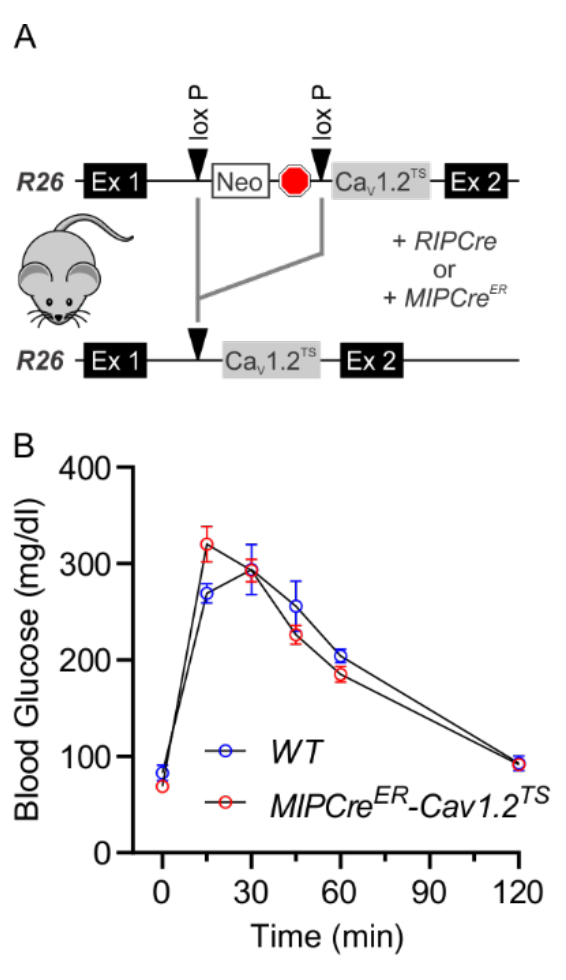
Ca_V_1.2^TS^ expression within beta cells is not sufficient to affect blood glucose. *A*, Schematic for transgenic Ca_V_1.2^TS^ expression from *Rosa26*. *B*, GTT in mice expressing Ca_V_1.2^TS^ under control of a tamoxifen inducible *Cre* recombinase restricted to beta cells driven by the mouse insulin promoter (or WT controls, n=3 each).

Although we assumed that glucose wasting was unlikely given the low-normal blood glucose and the lack of reported roles for Ca_V_1.2 in renal glucose reabsorption, we formally tested whether glycosuria contributed to hypoglycemia in the *TS2* mice. Unexpectedly, *TS2* mice displayed marked glycosuria (**Fig. 5A**) despite normal renal function (**Fig. 5B**). In the setting of normal renal function and low-normal baseline blood glucose, glycosuria suggests a defect in glucose reabsorption from the proximal tubule, which occurs in type 2 renal tubular acidosis (Fanconi syndrome). Accompanying the defect in glucose absorption in type 2 renal tubular acidosis is acidemia, which the *TS2* mice also displayed, as indicated by reduced serum bicarbonate compared to *WT* (**Fig. 5C**). Also similar to subjects with type 2 renal tubular acidosis who typically display polydipsia, the *TS2* mice drank more water (**Fig. 5D**). How the mutant Ca_V_1.2 channel contributed to glycosuria was not clear since Ca_V_1.2 protein was previously reported to be restricted to the inner medullary collecting ducts, and not found in proximal tubules (25). We therefore analyzed a recent human single cell RNA sequencing dataset of renal cells obtained by laser capture (26) and noted significant *CACNA1C* expression in tubules (**Fig. S4**). Further, we exploited a Ca_V_1.2 reporter mouse (*Ca_V_1.2^+/lacZ^*), in which *lacZ* (with a nuclear localization signal) disrupts one allele of *Cacna1c* and thus identifies Ca_V_1.2-expressing cells (27), and detected β-galactosidase signal in the proximal and distal convoluted tubules, as well as the expected intense signal within the renal vasculature and in the collecting ducts (**Fig. 5E-G**). While we do not expect that glycosuria contributes significantly to hypoglycemia in TS subjects, these data emphasize the widespread expression of Ca_V_1.2, even in non-excitable cells (3), and that physiological contributions of Ca_V_1.2 remain incompletely cataloged.

**Figure 5:**
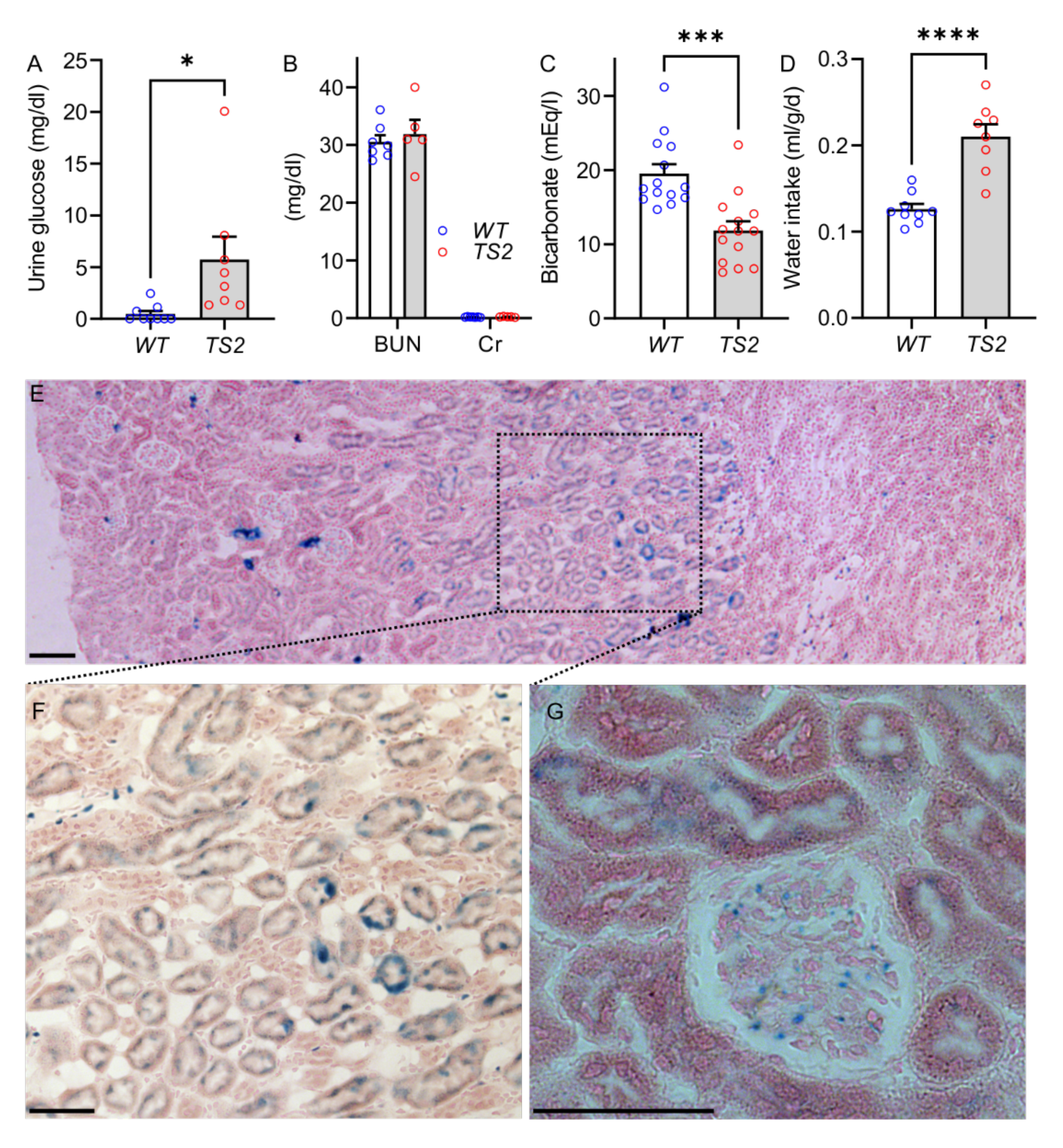
*TS2* mice display glycosuria. *A*, Urine glucose collected from fed mice (n=8-9, *t*-test, *, *p*<0.05). *B*, BUN and creatinine (n=5-7). *C*, Serum bicarbonate (n=14 each, *t*-test, ***, *p*<0.001). *D*, Water intake (normalized to body mass) during metabolic cage housing (n=8-9, *t*-test, ***, *p*<0.001). *E*, Histology and LacZ staining (with eosin counterstain) of kidney from a Ca_V_1.2 reporter mouse. *F*, Inset from E showing Ca_V_1.2 in the collecting ducts. *G*, Magnified image of a glomerulus and proximal tubules. Scale bars, 50 µm.

Because of the broad expression of Ca_V_1.2 in endocrine tissue (3), we investigated whether deficient counterregulatory hormone responses contributed to hypoglycemia in the *TS2* mice. An initial indication of blunted counterregulatory responses arose from the abnormally slow recovery of blood glucose levels in *TS2* mice after insulin administration. In an ITT of mice fasted for 2 h, we saw the expected initial drop in blood glucose followed by the subsequent initiation of recovery towards normoglycemia in *WT* mice. Blood glucose continued to decrease in *TS2* mice, however, despite lowering insulin to 1 unit/kg for the ITT, suggesting enhanced insulin sensitivity and/or deficient counterregulatory response to hypoglycemia (**Fig. 6A**).

**Figure 6:**
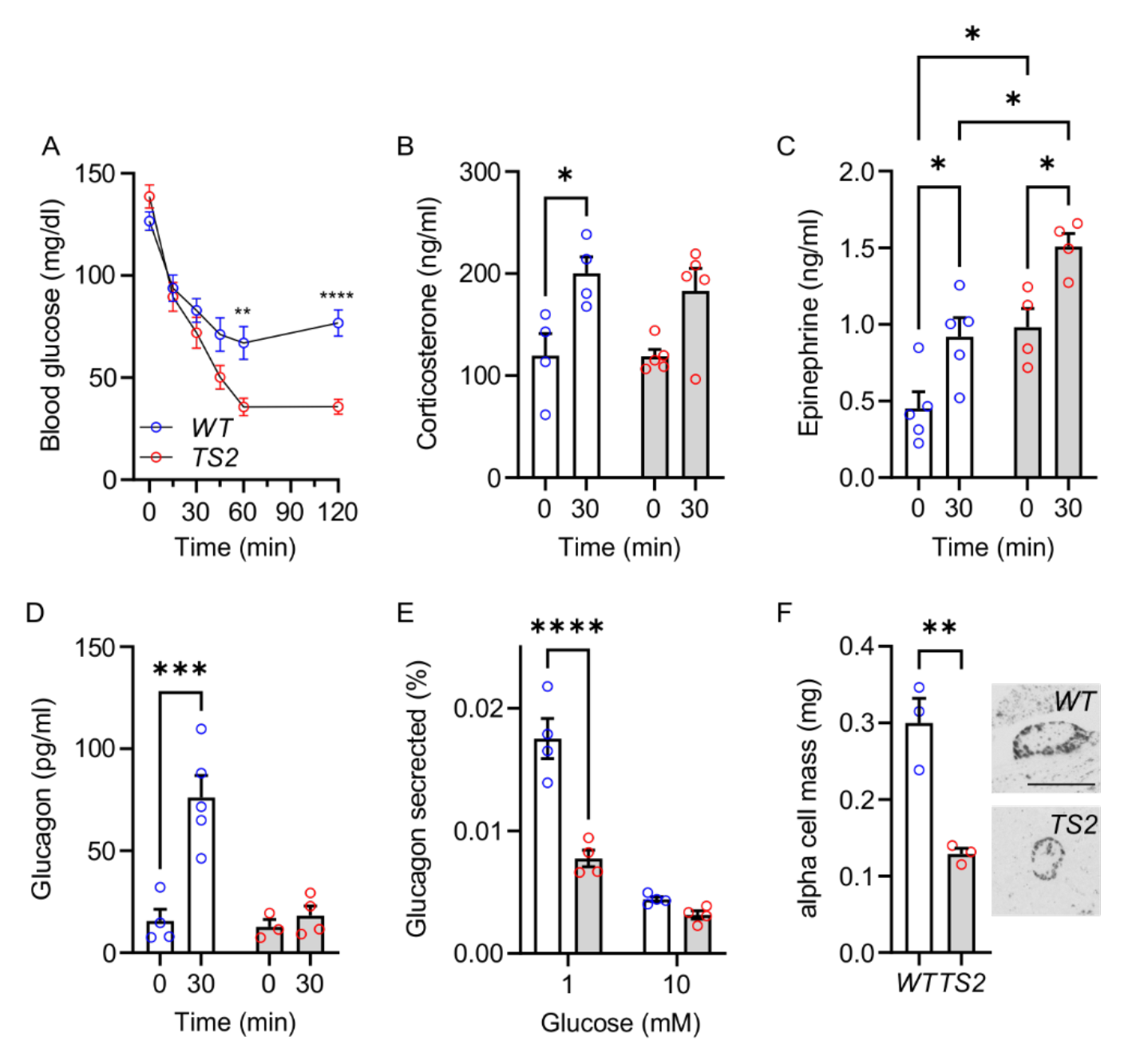
*TS2* mice display a blunted counterregulatory response. *A*, Insulin tolerance test (n=16-19, ANOVA corrected for multiple comparisons, **, *p*<0.01; ****, *p*<0.0001). *B-D*, Serum corticosterone (*B*), epinephrine (*C*), and glucagon (*D*) before (0) and 30 min after insulin administration (n=3-5, ANOVA, corrected for multiple comparisons, *, *p*<0.05; **, *p*<0.01; ***, *p*<0.001). *E*, Glucose regulated glucagon secretion from isolated islets (n=4 mice each, ANOVA, corrected for multiple comparisons, ****, *p*<0.0001). *F*, Pancreatic α cell mass (n=3 mice/genotype; n=3-4 sections/mouse). Inset shows exemplar α cell staining. Scale bar, 200 µm.

We considered that more efficient glucose utilization could contribute to the larger drop in blood glucose in response to insulin in the *TS2* mice, so we analyzed by immunoblot the relative amount of phosphorylation of Akt at S573 (pAkt S573) or phosphorylation of AMPK, downstream targets of insulin signaling, in lysates from skeletal muscle mice obtained in the fed state. No differences in insulin-induced normalized pAKT/AKT or pAMPK/AMPK ratios were detected between genotypes (**Fig. S5**). Thus, we concluded that a major contributor to the lower blood glucose in *TS2* mice was likely an insufficient counterregulatory response.

We therefore queried serum corticosterone, epinephrine, and glucagon, each of which showed in *WT* the expected increase in response to hypoglycemia induced by insulin (**Fig. 6B-D**). Insulin administration also elicited corticosterone and epinephrine increases in *TS2* mice (**Fig. 6B-C**), suggesting that those two counterregulatory hormone responses remained intact. Epinephrine values in *TS2* mice, however, were elevated compared to *WT*, both at baseline and 30 min post insulin administration. The elevated epinephrine in *TS2* mice compared to *WT* was unexpected, since a previous study (15) of isolated chromaffin cells from TS2-neo mice suggested decreased catecholamine secretion (measured by capacitance). This apparent inconsistency could reflect the measurement of capacitance change in chromaffin cells in the previous study vs. quantification of serum epinephrine in this study or reflect differences in mouse models (CRISPR knockin here vs. retained Neo cassette and consequent effect on the mutant allele’s expression in the previous study). Alternatively, the discrepant results could reflect the difference between measuring effects in isolated cells (previous study) vs. *in vivo* serum levels (here), which are subject to additional regulation, as we observed for insulin (**Fig. 2C-D**). Nevertheless, these data suggest that while hypoinsulinemia is likely a major contributor to the reduced adipose depots (**Fig. 3A-B**), elevated catecholamines and consequent increased lipolysis may be an additional contributor.

Although the corticosterone and epinephrine counterregulatory responses were intact, glucagon did not rise in response to insulin-induced hypoglycemia in the *TS2* mice (**Fig. 6D**), suggesting failure of that key counterregulatory hormone response. We further evaluated glucagon secretion and found that low glucose (1 mM) elicited 56% lower glucagon secretion in isolated islets from *TS2* compared to WT mice (**Fig. 6E**). Moreover, the alpha cell mass in *TS2* mice was reduced compared to *WT* (**Fig. 6F**). These data demonstrate deficiencies in pancreatic alpha cell mass and function in TS2.

The apparent discrepancy between the low insulin measured in *TS2* (vs. WT) serum and the lack of a genotype difference in insulin secretion from isolated islets (compare **Fig. 2C** and **Fig. 2D**) prompted us to consider whether glucose sensing and consequent regulation *in vivo* was perturbed. Specifically, since various hypothalamic nuclei control multiple aspects of glucose homeostasis and Ca_V_1.2 is broadly expressed in neurons, we hypothesized that Ca_V_1.2^TS^ expressed in the hypothalamus contributed to hypoglycemia in *TS2* mice by affecting glucose sensing. Using the Ca_V_1.2 reporter mouse, we queried where within the hypothalamus Ca_V_1.2 was enriched. **Fig. 7A** shows prominent lacZ signal in the arcuate nucleus and the paraventricular hypothalamus. Here, we focused specifically on the arcuate nucleus as POMC-expressing neurons within the arcuate nucleus are glucose sensing and regulate glucose homeostasis; manipulation of these neurons affects serum glucose (28, 29). Additionally, since *Pomc^+^* neurons show unusual step-like and long-lasting changes in their membrane potential in response to synaptic input due to persistent Na^+^ current through Na_V_1.7 channels (30), we hypothesized that these neurons are well tuned to respond to consequences of prolonged Ca^2+^ influx through Ca_V_1.2^TS^ mutant channels with their inherent defective voltage-dependent inactivation properties. We therefore focused on whether Ca_V_1.2^TS^ in *Pomc*^+^ neurons affected glucose sensing. We used the floxed-STOP TS mutant Ca_V_1.2 (Ca_V_1.2^TS^) line or its floxed-STOP wild-type Ca_V_1.2 (Ca_V_1.2^WT^) line as a control and activated the respective transgenes specifically in *Pomc*^+^ neurons by crossing the mice to a line that expressed *Cre* recombinase driven by the *Pomc* promoter (31) (**Fig. 7B**). In a GTT with mice expressing *Ca_V_1.2^TS^* specifically in *Pomc*^+^ neurons, we observed a reduction in blood glucose excursion similar to *TS2*, suggesting that expression of Ca_V_1.2^TS^ solely in *Pomc*^+^ neurons is sufficient to perturb glucose sensing (**Fig. 7C**). Notably, expressing the Ca_V_1.2^WT^ channel specifically in *Pomc*^+^ neurons had no effect on blood glucose (**Fig. 7D**) during a GTT, suggesting that the observed effects of Ca_V_1.2^TS^ in *Pomc*^+^ neurons derived from the Ca_V_1.2^TS^ mutant channel and not simply from transgenic Ca_V_1.2 overexpression. Although the difference between the Ca_V_1.2^WT^ and Ca_V_1.2^TS^ transgenes is the single nucleotide conferring the G406R mutation!thus unlikely to affect expression!we confirmed experimentally that the lack of an effect with the Ca_V_1.2^WT^ transgene was not due to reduced or absent expression of Ca_V_1.2^WT^ by measuring transgene protein by immunoblot. To obtain sufficient material (since *Pomc^+^* neurons are few in number), we analyzed expression of Ca_V_1.2^WT^ and Ca_V_1.2^TS^ in the heart (using the *MerCreMer* driver as in **Fig. S3B-C**) and we observed similar Ca_V_1.2 expression in the transgenic animals expressing Ca_V_1.2^WT^ or Ca_V_1.2^TS^ (**Fig. S6**).

**Figure 7:**
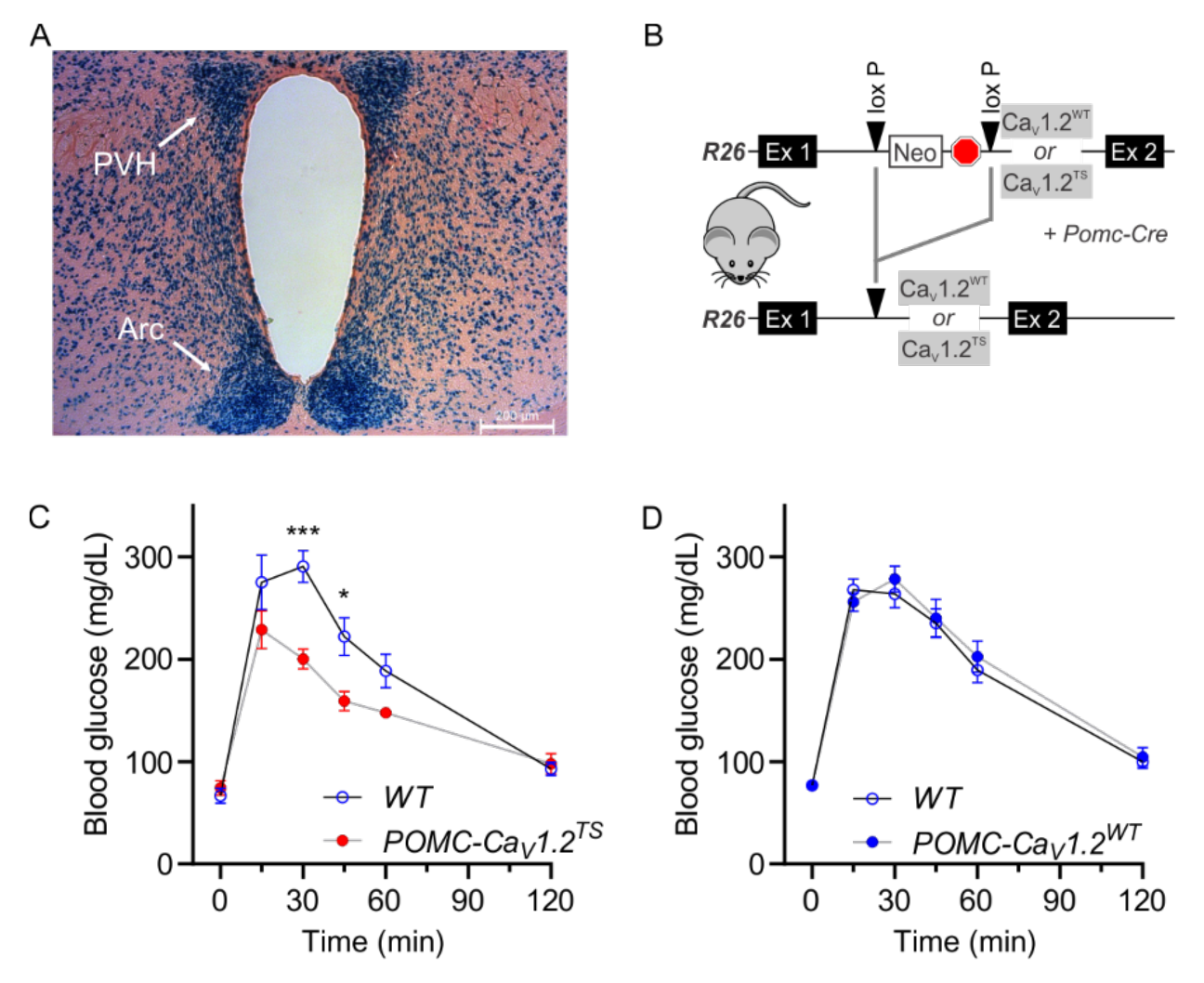
Expression of Ca_V_1.2^TS^ in *Pomc^+^* neurons affects glucose sensing. *A*, Histology and LacZ staining (with eosin counterstain) of coronal section through the hypothalamus from a Ca_V_1.2 reporter mouse. Arc, arcuate nucleus; PVH, paraventricular hypothalamic nucleus. *B*, Schematic for transgenic Ca_V_1.2^TS^ or Ca_V_1.2^WT^ expression restricted to *Pomc^+^* neurons from the *Rosa26* locus. *C*, GTT for mice expressing Ca_V_1.2^TS^ in *Pomc^+^* neurons or controls (n=4-5, ANOVA corrected for multiple comparisons, *, *p*<0.05; ***, *p*<0.001). *D*, GTT for mice expressing Ca_V_1.2^WT^ in *Pomc^+^* neurons or controls (n=8, each).

Altering POMC neuron excitability can affect food intake and body weight (30). Although we do not expect that the presence of a Ca_V_1.2^TS^ channel would alter neuronal excitability in POMC neurons that have Na^+^ channel-driven action potentials, but rather alter the neuron"s response by affecting the resultant Ca^2+^ influx through inactivation-deficient Ca_V_1.2^TS^ channels, we analyzed body mass in the *POMC-Ca_V_1.2^TS^* mice compared to their *WT* littermate controls. We observed no difference between genotypes (**Fig. S7**). Thus, the reduced weight and reduced food intake in the *TS2* mice (**Fig. S1B-C**) likely does not arise from POMC neuron expression of the Ca_V_1.2^TS^ channel and further emphasizes the multifactorial contributions to the specific phenotypes.

## Discussion

Together, our data demonstrate that the proposed hyperinsulinism driven by expression of the mutant Ca_V_1.2^TS^ channel in pancreatic beta cells is not the etiology for the severe episodic hypoglycemia suffered by TS patients. Several independent lines of evidence bolster this conclusion. First, we show that serum insulin levels are reduced, not elevated, in the *TS2* mice. This reduction in serum insulin occurs even though pancreatic beta cells in islets isolated from *TS2* mice show a normal response to glucose. Second, we show that Ca_V_1.2^TS^ channels, even when expressed specifically in pancreatic beta cells, are not sufficient to drive increased insulin secretion. This transgenic approach exploiting a cDNA demonstrates that the G406R mutation is incapable of increasing insulin secretion, independent of inclusion of exon 8 or exon 8A. These data are strengthened by the use of two different beta cell-specific promoters. The failure of these transgenes to lower blood glucose cannot be attributed to transgene inefficacy, as we showed efficacy in cardiomyocytes (**Fig. S3**) and in previous mouse models testing Ca_V_1.2^TS^ in bone physiology (32-34). Rather, we suggest that the failure of Ca_V_1.2 channels to drive increased insulin secretion in pancreatic beta cells points to the critical role of Ca_V_1.3 rather than Ca_V_1.2, as shown in a recent single cell sequencing data set (35) (**Fig 4C**). This is consistent with data from humans (36). A role for Ca_V_1.3 rather than Ca_V_1.2 is logical when considering that the resting membrane potential of beta cells is ∼ −70 mV (37), thus positioned to rely more upon Ca_V_1.3 for which the V_1/2_ for activation is significantly more negative than for Ca_V_1.2 (38).

These data contrast to a recent report of a *CACNA1C* gain of function mutation (L566P) discovered in a patient with congenital hypoglycemia (12). However, no accompanying data definitively link that mutation as causal. Further, the mutated residue is encoded by an invariant exon, so all *CACNA1C* transcripts from the mutant allele should contain the gain of function mutation (characterized only in a heterologous system), including those expressed in the heart. Yet the subject did not display a prolonged electrocardiographic QT interval nor any other phenotypes attributed to the canonical gain of function *CACNA1C* mutations in TS. Whether any *CACNA1C* mutations contribute to hyperinsulinism, therefore, remains unestablished. Further supporting our hypothesis that TS patients do not suffer from hyperinsulinism is that at least two TS subject were noted to have urinary ketones in the setting of hypoglycemia (12), a physiology inconsistent with a hyperinsulinism state. We also observed elevated serum ketones in the *TS2* mice. In summary, our data here with a knockin TS model and with transgenic Ca_V_1.2^TS^ channels, allow the approach for TS patients to be refocused for more productive therapeutic interventions.

Consistent with the many phenotypes exhibited by TS patients (1, 2) and coupled with the broad expression of *CACNA1C* and the similarly broad physiological effects of Ca_V_1.2, some of which are still not well understood nor explored (3), our data identify several alternative mechanisms for hypoglycemia in TS. Most prominently, we observe a perturbed counterregulatory response marked by intrinsic alpha cell dysfunction with reduced alpha cell mass and increased epinephrine. The mechanisms underlying alpha cell dysfunction are unknown, but may not involve expression of the mutant Ca_V_1.2 channel in alpha cells, since single cell sequencing data (36) suggest that the dominant L-type Ca^2+^ channel is Ca_V_1.3, not Ca_V_1.2 (**Fig S8**). In contrast to the absence of an apparent direct effect of the mutant Ca_V_1.2 channels in alpha cells, the abnormal baseline serum epinephrine in *TS2* mice likely arise from increased Ca^2+^ influx through Ca_V_1.2 in adrenal chromaffin cells driving increased epinephrine release. A role for Ca_V_1.2 in adrenal chromaffin cells has been established with the TS-neo mouse, in which decreased epinephrine release from isolated chromaffin cells was demonstrated (15). The reason for the opposing result in that mouse model is not clear, but the contrast of reduced epinephrine from isolated cells (TS- neo) compared to elevated epinephrine measured in serum (*TS2*) are analogous to the contrast between low serum insulin and normal insulin secretion from isolated islets we report here and are indicative of the importance of measuring the integrated physiologic response.

Further, we capitalized on the apparent inconsistency between the low serum insulin levels in *TS2* mice yet normal beta cell function in isolated islets to uncover a glucose regulatory role for Ca_V_1.2 within POMC expressing neurons in the arcuate nucleus. POMC neurons are glucose excited, increasing their firing rate in response to elevation in extracellular glucose (28), yet how POMC neurons affect glucose homeostasis is complex and remains incompletely understood. Insight is further complicated by data showing that POMC neurons are inhomogeneous in their cellular properties and downstream consequences (39). Expression of a mutant Kir6.2 K^+^ channel subunit, a component of the ATP sensitive K_ATP_ K^+^ channel, in POMC neurons, which prevents depolarization of baseline membrane potential, blocks the glucose-stimulated increase in firing rate and increases blood glucose during a GTT (28). Yet chemogenetic silencing of POMC neurons with a designer receptor exclusively activated by a designer drug (DREADD) decreases baseline blood glucose while having no effect on blood glucose during a GTT (29). Interestingly, chemogenetic activation of POMC neurons also does not affect blood glucose (29, 40), perhaps suggesting a critical threshold for Ca^2+^ influx. Indeed, the step-like changes in POMC neuron membrane potential in response to discrete synaptic inputs (30) are perfectly poised to trigger sustained openings of the inactivation-defective Ca_V_1.2^TS^ channels but not the inactivation-competent Ca_V_1.2^WT^ channels, providing a rationale why overexpression of the Ca_V_1.2^WT^ transgene did not affect blood glucose in our experiments. Thus, channel kinetics, not just increased Ca^2+^ influx, likely underlie the observed effects. Defining the specific cellular consequences of Ca_V_1.2^TS^ expression on POMC neurons and the resultant neural circuits affected awaits in-depth characterization of electrical and Ca^2+^ handling properties of those neurons as well as defining further how those neurons, when burdened with a Ca_V_1.2 channel with aberrant inactivation, contribute to glucose homeostasis.

Unexpectedly, glycosuria also appears to contribute to hypoglycemia in *TS2* mice. No data are available about glycosuria in TS patients. Thus, 2hile glycosuria is unlikely a major contributor to hypoglycemia, this finding underscores the utility of the animal models and the comprehensive physiological investigations we employed, allowing us to define novel physiological roles for Ca_V_1.2. How a mutant Ca_V_1.2 channel would cause type 2 renal tubular acidosis is unclear. *CACNA1C* has not previously been associated with type 2 renal tubular acidosis, perhaps due to the rarity of viable subjects with *CACNA1C* mutations and the severity of other life-threatening phenotypes. Expression and function of Ca_V_1.2 in the proximal tubule have not been definitively studied, thus providing motivation for future studies.

Finally, because of the multiple *Cacna1c*-dependent causes driving hypoglycemia in *TS2* mice, it is interesting to speculate why hypoglycemia in TS patients is episodic and not completely penetrant. For this, we consider the rarity of *CACNA1C* mutations: With a probability of being loss-of-function intolerant (pLI) score = 1 and the observed / expected (oe) metric = 0.05 (41), *CACNA1C* is predicted to be highly intolerant of mutations. Consistent with that hypothesis, TS is rarely inherited and, when inherited the parent may be an unaffected or barely affected mosaic for the *CACNA1C* mutation (42). We thus further speculate that many, if not most, TS subjects are mosaic for the *CACNA1C* mutation. The degree of mosaicism and the specific tissues in which the mutant allele is present determines the presence and severity of specific TS-associated phenotypes, such as hypoglycemia. With the strong physiologic drive towards homeostasis, we speculate that a complex phenotype like hypoglycemia may manifest only in subjects with a sufficient mosaic mutational load to perturb the multiple systems and cause hypoglycemia. In this context, the utility of a mouse model provides an invaluable resource.

## Methods

### Animals

*C57BL/6J* mice (Stock #00064), *Ca_V_1.2^+/lacZ^* reporter mice (Stock #0005783), *POMC-Cre* (Stock #005965), *MIP-Cre^ER^* (tamoxifen-inducible mouse insulin promoter, Stock #024709), and *RIP-Cre* (rat insulin promoter, Stock #003573) were purchased from Jackson Laboratory. STOP-floxed *Ca_V_1.2^WT^* or a G406R mutant (*Ca_V_1.2^TS^*) mice were described previously (21). Timothy syndrome (*TS2*) mice were generated with CRISPR/Cas9 technique by Duke University Transgenic Core Facility using 5’ ctggttctcggtgttttgagCGG 3’ (capital letters = PAM sequence) for the guide RNA and 5’ tcatctttggatcctttttcgttctaaatctggttctcggtgttttgagc**A**ggtaagctgaccacttctgtgtcctctttacaacgcagccgagcaaggtcc caggttcaatgcagaatcctcagagacaggaccctaacaggccc 3’ (capital letter = point mutation) for the repair oligo. *TS2* mice were backcrossed more than eight times to *C57BL/6J*. Except when food restricted, mice received a normal chow diet. Because of known variability of female mice in metabolic studies (43), analyses were restricted to male mice except where indicated. Cre-mediated recombination was achieved by intraperitoneal injection of 75 μg/g body weight of tamoxifen dissolved in corn oil for 5 consecutive days.

### Electrocardiograms (ECGs)

Three-lead surface electrocardiograms were recorded on anesthetized animals (Avertin, 250mg/kg i.p.) using a PowerLab (ADInstruments) recorder and were analyzed with Labchart Pro software (ADInstruments). QT intervals were obtained automatically and confirmed by visual inspection. For ECGs after adrenergic stimulation, anesthetized mice received isoproterenol (1.5 mg/kg) via i.p. injection.

### Glucose and pyruvate tolerance tests (GTT)

Mice (8-20 weeks old; age indicated in text for each experiment) were food restricted for 16 hours, with water ad libitum. Blood from the tail vein was used to measure glucose levels using a glucometer (Abbott) immediately before and 15, 30, 45, 60 and 120 min after intraperitoneal injection with 2 g/kg (body weight) glucose.

### Insulin tolerance test (ITT)

Mice (14-20 weeks old) were food restricted for 2 hours, with water ad libitum. Blood from the tail vein was used to measure glucose levels using a glucometer (Abbott) immediately before and 15, 30, 45, 60 and 120 min after intraperitoneal injection with 1.0 U/kg (body weight) insulin (Humulin R-100).

### X-gal staining

Mouse brain and kidney tissues were collected from *Ca_V_1.2^lacz/+^* reporter mice (14-20 weeks old), embedded in Tissue-Tek^®^ O.C.T. Compound (Sakura), and sectioned at 10 um. Sections were fixed with 0.2% glutaraldehyde and 2mM MgCl_2_ in phosphate buffered saline (PBS) and incubated in the X-gal staining solution (Gold Biotechnology) over night at 37 °C. Nuclear fast red (Vector Laboratories) was used for counterstaining.

### Pancreatic beta cell and α cell mass immunohistochemistry, imaging, and mass calculation

To determine beta cell and α cell mass, insulin or glucagon immunohistochemistry was performed in at least 3 non-consecutive pancreatic sections for each mouse (14-20 weeks old) as previously described (44). Briefly, anti-insulin (1:10,000, Abcam) or anti-glucagon (1:200, Cell Signaling) antibody was incubated overnight at 4°C, followed by incubation with biotinylated secondary antibody (1:500, Vector Laboratories). Then avidin–biotin peroxidase complex (Vectastain ABC kit, Vector Laboratories) was added for 30 min. Sections were developed using peroxidase substrate (Vectastain DAB kit, Vector Laboratories). Whole sections were scanned using a Keyence BZX800 whole-slide scanner. The fraction of insulin or glucagon-positive areas compared to total pancreatic tissue area was determined with Fiji/ImageJ (NIH, Bethesda, MD). beta cell and α cell mass were then determined by multiplying the obtained fraction by the initial pancreatic wet weight.

### Pancreatic islet isolation and static glucose stimulated insulin or glucagon secretion assays

Mouse pancreatic islets (14-20 weeks old mice) were isolated by perfusion of the pancreases with CiZyme (VitaCyte) through the common hepatic duct. Pancreases were removed and digested at 37 °C for 17 minutes. After two washes with RPMI medium with 3% FBS, islets were separated into a gradient using RPMI medium and Histopaque (Sigma-Aldrich). Islets were then hand-picked to avoid exocrine contamination and processed for different applications. For static glucose stimulated insulin secretion assays, we handpicked islets and placed them into basal Krebs buffer containing 3 mM, 11 mM, or 20 mM glucose for insulin secretion; and 1 mM or 10 mM glucose (both with 10 mM arginine supplementation) for glucagon secretion followed by transfer into Krebs solution containing the indicated concentrations of glucose. After 45 min incubation at 37°C, islets were pelleted, and supernatants were collected to measure secreted insulin. We lysed islet pellet with RIPA buffer to assess intracellular insulin or glucagon content. Insulin or glucagon was measured by ELISA (Mercodia or Chrystal Chem)

### Serum insulin, glucagon, leptin, corticosterone, epinephrine, liver glycogen store, and urine glucose measurements

For insulin analysis, blood was collected from 14-20 weeks old mice during GTT. Serum insulin was measured with mouse ultrasensitive insulin ELISA (Crystal Chem). For glucagon, leptin, corticosterone, and epinephrine, blood was collected from 14-20 weeks old mice during ITT. Serum glucagon was measured with mouse glucagon ELISA (Crystal Chem). Serum leptin was measured with mouse leptin ELISA (Crystal Chem). Serum corticosterone was measured with mouse corticosterone ELISA (Crystal Chem). Serum epinephrine was measured with epinephrine ELISA (ImmuSmol). Liver glycogen was measured with Glycogen Assay Kit II (Abcam) according to the manufacturer’s instructions. Urine was collected from animals at fed state, and urine glucose was measured with Mouse Glucose Assay Kit (Crystal Chem).

### Adipose tissue collection and analysis

Subcutaneous white adipose tissue (sWAT) and epididymal white adipose tissue (eWAT) were excised from WT or TS2 mice (14-20 weeks old) and photographed immediately after dissection for gross morphologic comparison. Each tissue was weighed and normalized to body weight.

### Smooth Muscle Isolation

Smooth muscle cells were isolated from the colon of adult TS2 mice. Isolation procedure was performed, as previously described (45), in low Ca^2+^ Tyrode solution containing (in mM) 137 NaCl, 2.7 KCl, 0.008 CaCl_2_, 0.88 MgCl_2_, 0.36 NaH_2_PO_4_, 12 NaHCO_3_ and 5.5 glucose. Mice were euthanized and the colon was isolated. Colon was pinned down and cut open following which the mucosal layer was scrapped off to expose the muscle layer. The muscle layer was cut into pieces and incubated in 5 ml of digestive solution containing 1.5 mg collagenase (Worthington), 1 mg trypsin (Sigma) and 5 mg BSA (Sigma) at 37°C. Tissue was then triturated using a glass pipette to dissociate cells and the solution was monitored under the microscope to check for isolated cells. The partially digested tissue was then transferred to an enzyme free solution to complete the isolation through trituration. Isolated cells were stored on ice and used for electrophysiology recordings for 6 h.

### β-hydroxybutyrate (ketone bodies) measurement and serum bicarbonate analysis

Blood from the tail vein of fed-state mice (14-20 weeks old) was used to measure β-hydroxybutyrate using Lactate Plus (Nova Biomedical GmBH). Serum bicarbonate, blood urea nitrogen (BUN), and creatine were analyzed by Weill Cornell Medicine Laboratory of Comparative Pathology Core using the spectrophotometry (Beckman Coulter, Clinical Chemistry Analyzer AU680).

### Insulin-stimulated Akt/ phospho-Akt and AMPK/ phospho-AMPK quantification

Insulin (1 unit/kg, Human R [Lilly]) diluted in PBS or PBS was administered i.p. 20 min before mice were euthanized and the quadriceps were quickly excised. Whole tissue lysate was prepared from the mouse liver tissues by homogenizing with The Bullet Blender Blue 24 (Scientific Instrument Services) in T-PER buffer (Thermo Fisher Scientific) containing Halt Protease and Phosphatase Inhibitor Cocktail (Thermo Fisher Scientific). Lysates were then rocked for 2 hours at 4 °C followed by centrifugation at 12,300 x g at 4 °C for 15 minutes and supernatants were collected after fat layer was removed. Protein concentration was quantified using a BCA Protein Assay Kit (Thermo Fisher Scientific). Approximately 10 µg of protein were loaded on Novex 8% Tris-Glycine gels (Invitrogen), electro-transferred to a polyvinylidene difluoride membrane (Thermo Fisher Scientific) for 60 minutes at 20 V. Membranes were blocked with 5% non-fat milk and 1% BSA in TBST for 1 hour at room temperature, and then incubated in primary antibodies; Pan-Akt (1:1000, Cell Signaling), pAkt (S437) (1:1000, Cell Signaling), AMPK (1:1000, Cell Signaling), pAMPK (1:1000, Cell Signaling), Vinculin, (Santa Cruz,sc-73614) overnight at 4 °C. Proteins were detected using SuperSignal West Pico Plus Chemiluminescence Substrate (Thermofisher, 34580) before imaging and quantification on a ChemiDocTM Touch Imaging System (Bio-Rad).

### Electrophysiology

Hippocampi from 1- to 2-day newborn *WT* and *TS2* mice were dissociated through enzymatic treatment with 0.25% trypsin and subsequent trituration. The cells were plated on glass coverslips previously coated with poly-D-lysine and laminin in 12-well cell culture plate. Hippocampal cells were grown in neurobasal A medium (ThermoFisher Scientific) supplemented with 2% B27, 2 mM glutamine, 10% heat-inactivated fetal bovine serum and 1% penicillin/streptomycin in 5% CO_2_ incubator at 37°C overnight and then this medium was replaced by one containing 2% B27, 0.5 mM glutamine, 1% heat-inactivated fetal bovine serum, 70 µM uridine and 25 µM 5-fluorodeoxyuridine.

Voltage-gated calcium current recordings were performed from hippocampal pyramidal shape neurons cultured for 15-16 days at room temperature. The current was induced with a 500 ms depolarization step at 0 mV from the holding potential of −60 mV using barium as the charge carrier and was sampled at 10 kHz and filtered at 2 kHz in the whole-cell voltage patch-clamp configuration with an Axopatch 200B amplifier (Molecular Devices). Peak and residual currents were obtained with Clampfit software (Molecular Devices), and mean current between 495 and 500 ms of the depolarization step was used as the residual current at 500 ms (r500).

The pipette solution for calcium current recording contained (in mM): 120 CsMeSO_3_, 5 tetraethylammonium chloride, 5 EGTA, 5 MgCl_2_, 2 Mg-ATP, 0.2 GTP, 5 Na_2_-phosphocreatine, and 5 HEPES, pH 7.3 with CsOH, osmolarity 290 mOsm/L; external solution contained (in mM): 110 NaCl, 5 KCl, 20 HEPES, 2 BaCl_2_, 2 MgCl_2_, 30 glucose, 0.0005 tetrodotoxin, 0.05 APV, 0.02 DNQX, 0.02 bicuculine, 20 tetraethylammonium chloride, and 2 4-aminopyridine, pH 7.3 with NaOH, Osmolarity 300-310 mOsm/L.

For smooth muscle cell electrophysiology, a HEKA EPC 10 amplifier was used for recording currents and Patchmaster software (HEKA) was used for pulse generation and data acquisition. Recordings were carried out at room temperature (22-25°C). Patch pipettes with 2-2.5 MΩ resistance were used for recordings (P97 – Sutter instrument). Pipettes were filled with solution containing (in mM) 100 L-aspartic acid, 30 cesium chloride, 1 MgCl_2_, 5 HEPES, 2 adenosine triphosphate (disodium salt), and 5 EGTA, with pH adjusted to 7.2 using cesium hydroxide. Cells were placed in bath solution containing (in mM) 135 NaCl, 5.4 KCl, 0.33 NaH_2_PO_4_, 5 HEPES, 1 MgCl_2_, 20 BaCl_2_, 5 D-glucose, with pH adjusted to 7.4 using NaOH. All chemicals were obtained from Sigma. Cell membrane was held at a resting voltage of - 90 mV in a whole cell patch clamp mode and stimulated to −5 mV for 150 ms to activate the Ca^2+^ channels.

### Food and water intake measurements

Food and water intake measurements were obtained from metabolic measurements acquired using the Promethion Metabolic Screening System (Sable Systems International, Las Vegas, NV) in metabolic cages maintained at a stable ambient temperature of 22.0 ± 1.0 °C with a 12-hour light/dark cycle by the Metabolic Phenotyping Center of Weill Cornell Medicine. Mice (14-20 weeks old) were single housed for the duration of the experiment and had ad libitum access to food and water. The mice were acclimated to the metabolic cages for 48 hours, followed by 24 hours of data recording. Food intake, water intake, and body mass were continuously monitored gravimetrically.

### RNA-Seq analysis

Data for mouse beta cells were analyzed as described (35). Data from human islets was obtained from Trischler *et al.* (36), and analyzed as described (35).

## Statistics

Statistical analyses were performed using GraphPad Prism, version 9.4.1. All averaged data are presented as mean ± SEM. Statistical significance was determined using 2-tailed *t* tests, 2-way ANOVA with post hoc correction for multiple comparisons, or *χ*^2^ test. *P* < 0.05 was considered statistically significant.

## Study approval

Animals were handled according to NIH Guide for the Care and Use of Laboratory Animals. The research was approved by the Weill Cornell Research Animal Resource Center (Protocol Number: 2016-0042).

## Acknowledgements/Sources of Funding

Supported by R01 HL151190 and R01 HD090132 (GSP) and R01 DK121140 and R01 DK121844 (JCL).

## Author Contributions

MM designed research studies, conducted experiments, acquired data, analyzed data, and wrote the manuscript. LEL, ID, AG, NG-B, H-GW, ARG, DSS EQW, ASB, KJ, AR-N, and AL conducted experiments, acquired data, and analyzed data. SOM, TEM, PT, KT, and JLC designed research studies and edited the manuscript. GSP designed research studies, analyzed data, and wrote the manuscript.

