## Supplemental Figures for "Multiple beta cell-independent mechanisms drive hypoglycemia in Timothy syndrome"

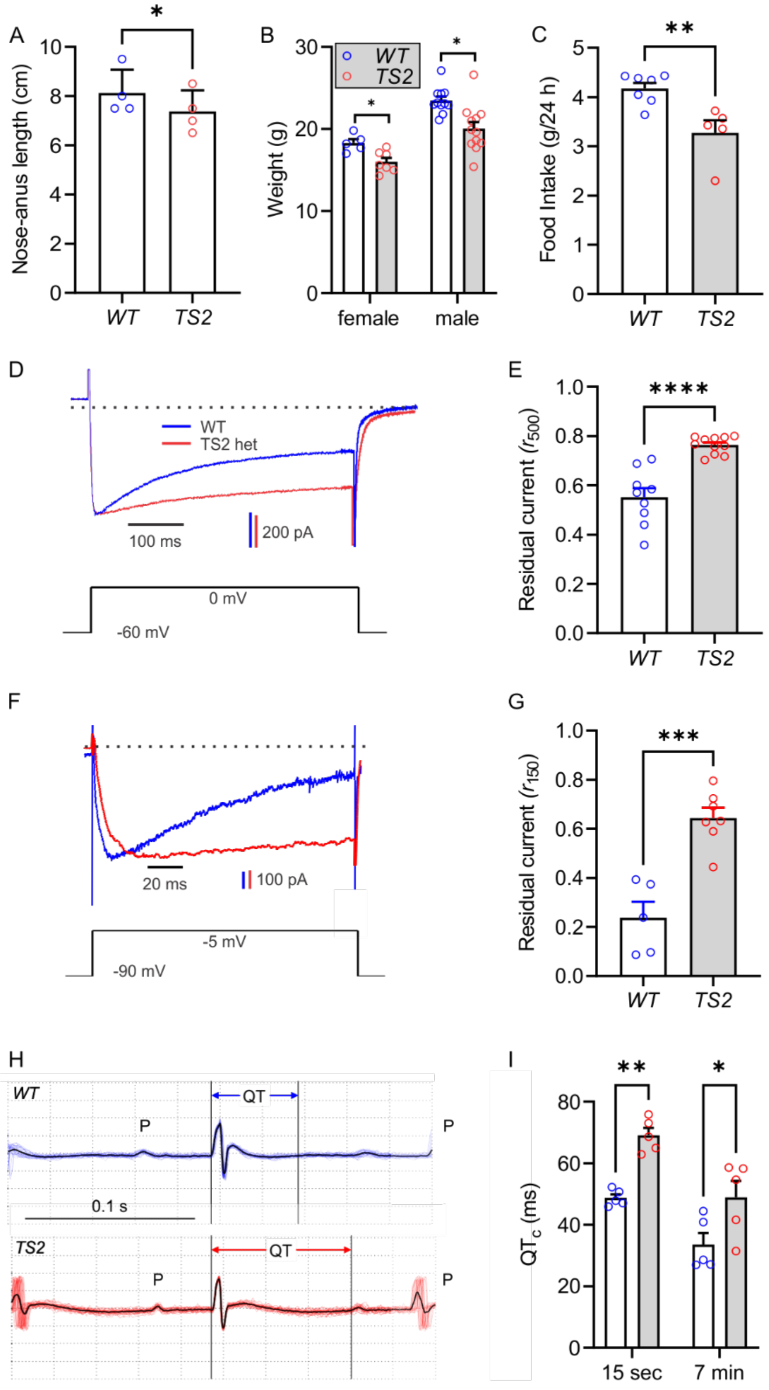


**Figure S1:** Phenotyping of the *TS2* mouse. *A*, Length (males, n=4 each genotype; *t*-test; *, *p*<0.05) and *B*, Weights (n=5-12; *t*-test; **, *p*<0.01, corrected for multiple comparisons) of *WT* and *TS2* mice. *C*, Food intake in metabolic cages (n=5-7, *p*<0.01, *t*-test). *D*, Exemplar $Ba^{2+}$ currents from isolated hippocampal neurons at 15-16 days *in vitro*. *E*, Residual ($r_{500}$) current at the end of a 500 ms test pulse to 0 mV (n=9-11; *t*-test; ****, *p*<0.001). *F*, Exemplar $Ba^{2+}$ currents from isolated colon smooth muscle cells. *G*, Residual ($r_{150}$) current at the end of a 150 ms test pulse to 0 mV (n=5-7; *t*-test; ****, *p*<0.001). *H*, Example ECGs. P waves and the QT interval are identified. *I*, Corrected QT interval (QTc) after isoproterenol administration (n=5 each; ANOVA, *, *p*<0.05; and **, *p*<0.01).

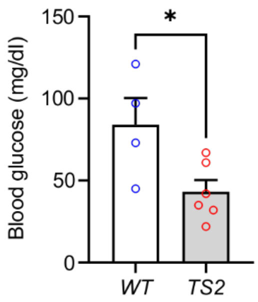


**Figure S2:** Blood glucose after 16 h fast in 8 week old mice (n=4-6; *t*-test, *, $p<0.05$).

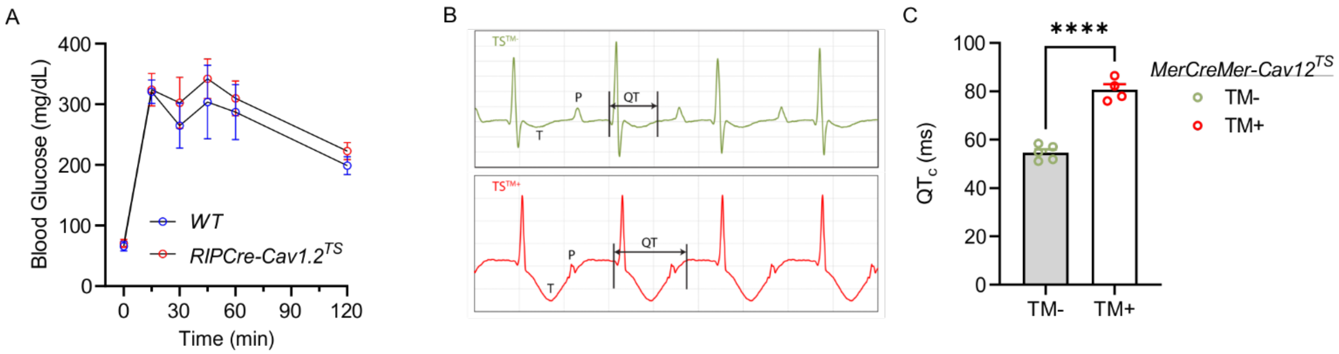


**Figure S3:** Controls for $Ca_V1.2^{TS}$ targeted to pancreatic β cells. *A*, GTT in mice expressing $Ca_V1.2^{TS}$ under control of a constitutive *Cre* recombinase restricted to β cells driven by the rat insulin promoter (or WT controls, n=3-4). *B*, Exemplar ECGs from mice expressing $Ca_V1.2^{TS}$ under control of a tamoxifen inducible *Cre* recombinase restricted to cardiomyocytes driven by the *MerCreMer* α-myosin heavy chain promoter before ($TS^{TM-}$) or 2 weeks after ($TS^{TM+}$) administration of tamoxifen. Example P and T waves, and the QT interval, are noted. *C*, The corrected QT interval (QTc) before or 2 weeks after administration of tamoxifen (n=4-5, *t*-test, ****, $p<0.0001$).

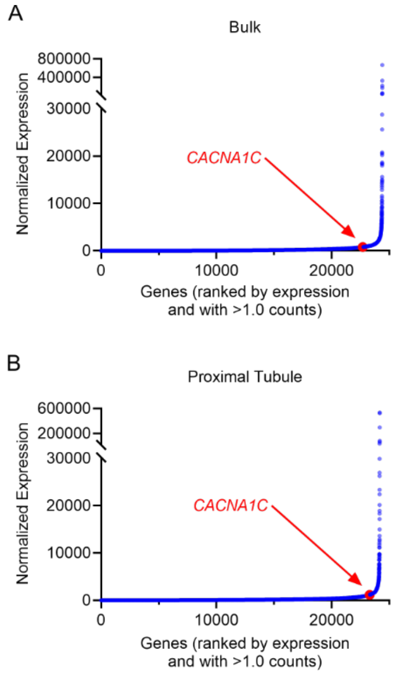


**Figure S4**: RNA sequencing data from Supplemental Table 5 in PMID: 35675394 (25). Data for bulk RNA sequencing data (*A*) and for proximal tubules (*B*) were averaged, sorted by average number of counts, and all genes with counts >1.0 were plotted. Expression of *CACNA1C* is highlighted in red.

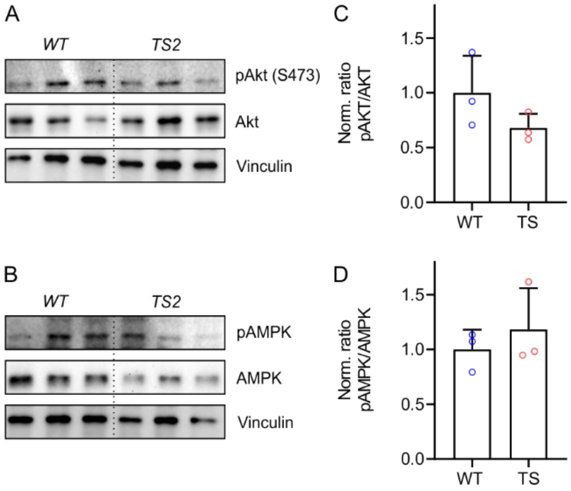


**Figure S5**: Post insulin receptor signaling in skeletal muscle (30 min after insulin administration) monitored by Akt phosphorylation at Ser 473 or AMPK phosphorylation. *A* and *B*, immunoblots for Akt (*A*) AMPK (*B*) each with vinculin immunoblots as a loading controls. *C*-*D*, Normalized band intensity for pAkt/AKT (*C*) and pAMPK/AMPK (*D*) ratios. n=3, each genotype.

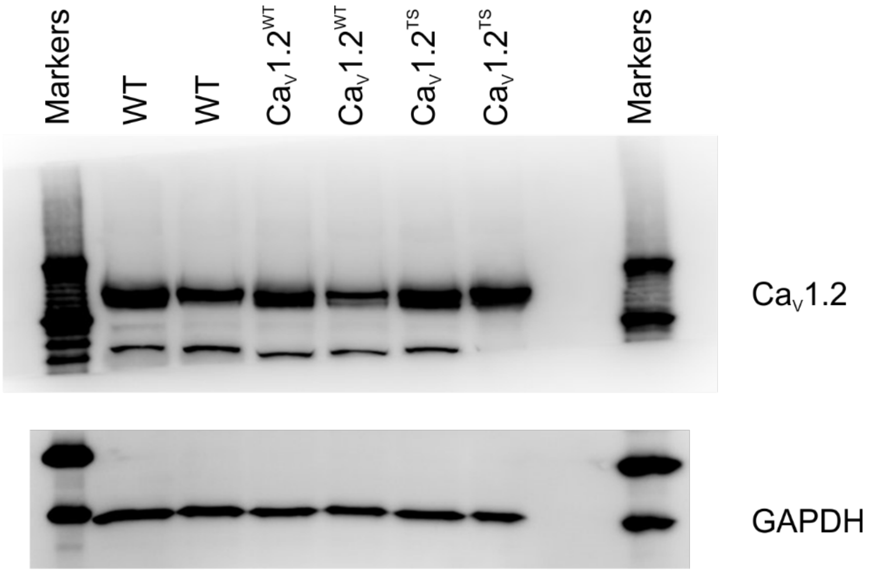


**Figure S6**: Expression of Ca$_V$1.2$^{TS}$ or Ca$_V$1.2$^{WT}$ in heart. Immunoblots for Ca$_V$1.2 or GAPDH expression in heart lysates from WT, or transgenic animals expressing either Ca$_V$1.2$^{TS}$ or Ca$_V$1.2$^{WT}$ driven by the *MerCreMer* promoter.

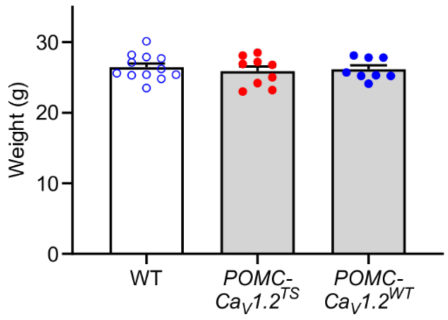


**Figure S7**: Weights of POMC transgenic animals and their WT littermate controls (n=8-12, *p*>0.05 one-way ANOVA).

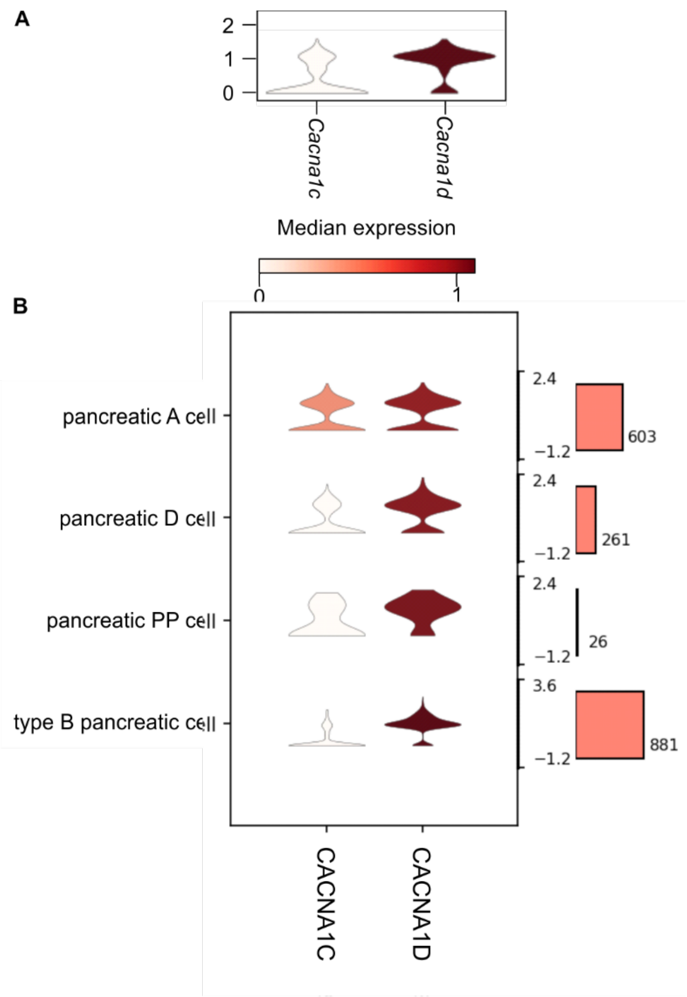


**Figure S8:** Expression of L-type $Ca^{2+}$ channel genes in mouse (A) and human (B) pancreatic islet cells form single cell RNA-seq data.
